# Rare SNP in the *HELB* gene interferes with RPA interaction and cellular function of HELB

**DOI:** 10.1101/2024.02.27.582415

**Authors:** Bertha Osei, Benjamin H. May, Clara M. Stiefel, Kirk L. West, Maroof Khan Zafar, Matthew D. Thompson, Erik Bergstrom, Justin W. Leung, Eric J. Enemark, Alicia K. Byrd

**Author notes:** Clara Stiefel, Department of Radiation Oncology, University of Texas Health Science Center San Antonio, San Antonio, Texas, 78229, USA.

## Abstract

HELB is a human helicase involved in initiation of DNA replication, the replication stress response, and regulation of double-strand DNA break repair. rs75770066 is a rare SNP in the HELB gene that affects age at natural menopause. rs75770066 results in a D506G substitution in an acidic patch within the 1A domain of the helicase that is known to interact with RPA. We found that this amino acid change dramatically impairs the cellular function of HELB. D506G-HELB exhibits impaired interaction with RPA, which likely results in the effects of rs75770066 as this reduces recruitment of HELB to sites of DNA damage. Reduced recruitment of D506G-HELB to double-strand DNA breaks and the concomitant increase in homologous recombination likely alters the levels of meiotic recombination, which affects the viability of gametes. Because menopause occurs when oocyte levels drop below a minimum threshold, altered repair of meiotic double-stranded DNA breaks has the potential to directly affect the age at natural menopause.

## INTRODUCTION

In addition to the loss of fertility, the decrease in estrogen levels after menopause leads to changes in disease risk in postmenopausal women (1). Incidence of cardiovascular disease increases two-fold after menopause (2, 3) and bone mineral density, which is inversely related to fracture risk, decreases ∼10% during the menopause transition and ∼1% annually in postmenopausal women (4, 5). On the other hand, risks of some cancers increase in women with later menopause, with women in the highest quintile of age at menopause (ANM) having a 15% increased incidence of estrogen receptor positive breast cancer and a 10% increase in estrogen receptor negative breast cancer (6). Additionally, the risks of uterine and ovarian cancer increase by 42% in women who enter menopause after age 55 (7).

Because menopause occurs when the oocyte pool is depleted to the point that it is insufficient to support ovulation (8, 9), factors affecting the size and quality of the oocyte pool are directly related to ANM. Recent genome wide association studies (GWAS) have illustrated that factors affecting genome stability play critical roles in regulation of reproductive senescence (6, 10–12). Two-thirds of the genomic regions containing ANM associated single-nucleotide polymorphisms (SNPs) are DNA damage response (DDR) genes (6, 10).

Multiple SNPs in the gene encoding the HELB helicase have been independently linked to ANM (6, 12–14). Of particular interest is rs75770066, a low-frequency, nonsynonymous SNP in *HELB*. Multiple reports link rs75770066 to an increase in ANM, but a decrease in ANM has also been reported (6, 12–14). These conflicting results are explained by a heterozygote effect—a single copy of rs75770066 increases ANM by ∼1 year while women homozygous for the rare allele have an ANM ∼1 year earlier (12). Because rs75770066 is a low-frequency variant (minor allele frequency = 3.6%), less than 0.2% of women are homozygous for the rare allele, while 3.5% of women are heterozygous.

Because multiple SNPs in HELB are independently associated with ANM, this suggests that HELB is a critical component of a process affecting reproductive senescence, but the mechanism of this relationship is unclear. HELB is a superfamily 1B (SF1B) DNA helicase in vertebrates (15, 16) that unwinds DNA with a 5′-to-3′ polarity (17). HELB is important for timely entry into S phase (17, 18), possibly by promoting formation of the pre-initiation complex (18). HELB also enhances survival of cells from DNA replication stress (19), and regulates homologous recombination (HR) for repair of double-stranded DNA breaks (DSBs) (20, 21). HELB interacts with the single-stranded DNA binding protein, RPA (19, 21), and this interaction is essential for HELB recruitment to laser induced micro-irradiation (21). HELB can also remove RPA filaments from single-stranded DNA (ssDNA) (22). The relationship between these cellular functions of HELB and alterations in ANM caused by rs75770066 is unknown.

HELB contains an N-terminal domain that is involved in protein-protein interactions and may bind to DNA (18, 22), a central helicase domain, and a C-terminal subcellular localization domain (23) (**Figure 1A**). rs75770066 results in a D506G substitution in HELB. D506 is located in an insertion (amino acids 496-526) within the helicase domain between helicase motifs I and Ia that is conserved in HELB family members but lacking in other Superfamily 1B helicases (**Figure 1A-B** and **Supplementary Figure S1**) (15). This HELB specific motif (HSM) is a similar region to the acidic motif (amino acids 493-517) previously identified by the Fanning lab (19) and is predicted by AlphaFold to extend from the leading edge of the 1A domain of the helicase (**Figure 1C**) (24, 25). This region contains 3 residues that interact with RPA: E499, D506, and D510 (19). Substitution of all three residues with alanine (E499A/D506A/D510A, or 3xA) interferes with HELB interaction with RPA (19) and reduces HELB localization to sites of laser-induced microirradiation (21). Consistent with biochemical data, AlphaFold predicts that the HSM interacts with the RPA70 N-terminal domain (NTD) (**Figure 1D**) (24). A recent structure of a peptide containing a portion of the HSM containing E499, D506, and D510 and the RPA70 NTD confirms the interaction of these residues with RPA (**Figure 1E**) (26). To gain insight into the mechanism by which rs75770066 alters ANM, we measured the effect of a D506G substitution in HELB on helicase activity, protein-protein interactions, and the cellular activities of HELB.

**Figure 1.**
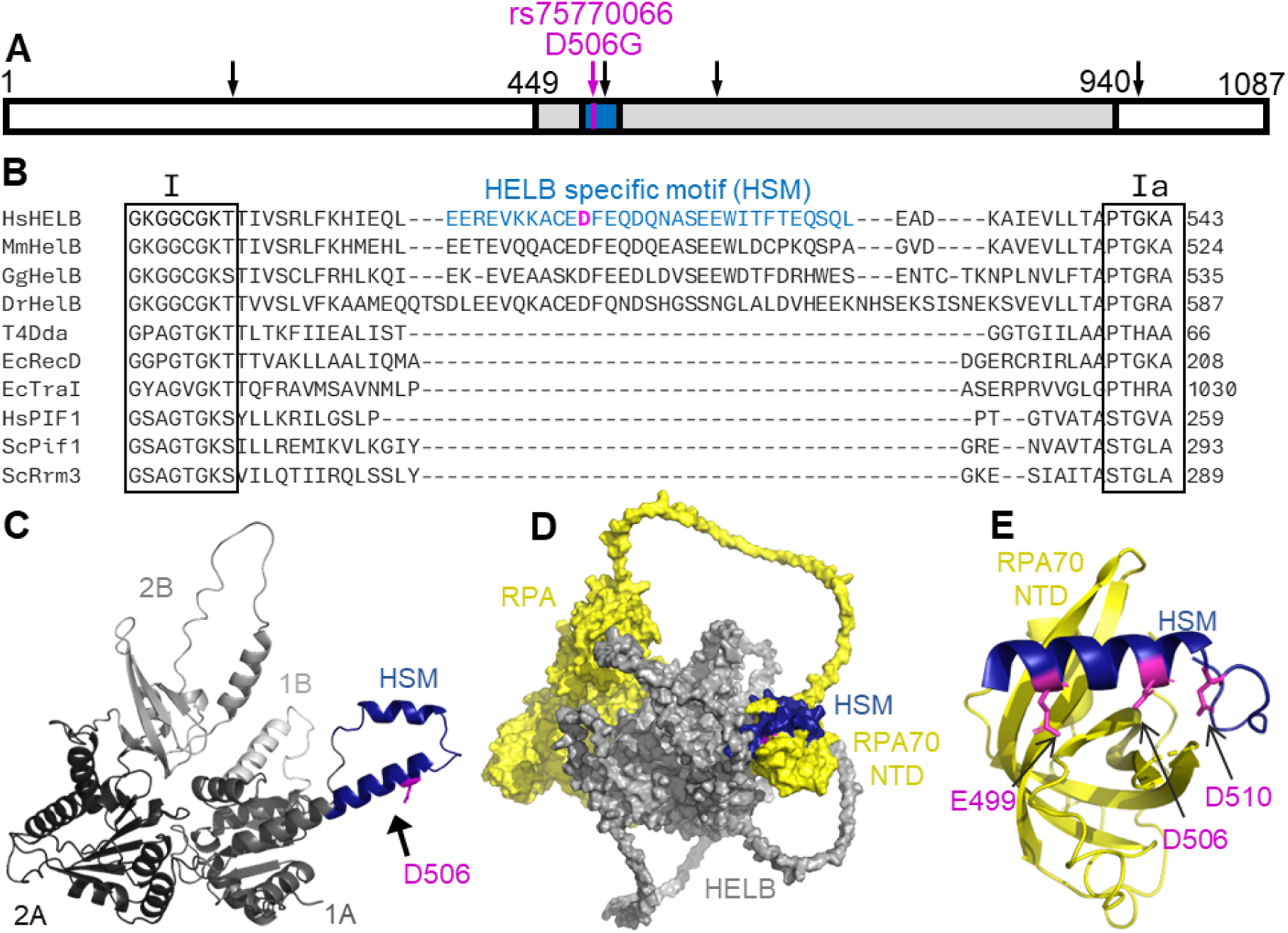
HELB specific motif is an extension of the 1A helicase domain that interacts with RPA. (**A**) Schematic of the HELB protein. The helicase domain is shaded with the HELB specific motif in blue. Locations of amino acid substitutions due to SNPs associated with age at natural menopause are marked with arrows. (**B**) Sequence alignments of *Homo sapiens* HELB, *Mus musculus* HelB, *Gallus gallus* HelB, *Danio rerio* HelB, Bacteriophage T4 Dda, *E. coli* RecD, *E. coli* TraI, *Homo sapiens* PIF1, *Saccharomyces cerevisiae* Pif1, and *Saccharomyces cerevisiae* Rrm3 helicases by Clustal Omega shows a HELB specific motif (HSM) (blue) exists between helicase motifs I and Ia. D506 (magenta) is located within this HSM. (**C**) AlphaFold structure prediction of HELB helicase domain (gray) illustrates the HSM (blue) is an extension of the helicase domain. D506G (magenta) is located in the HSM. The 1A and 2A conserved RecA-like domains are colored in dark gray and black, respectively. The predicted strand separation wedge (1B) is colored in light gray, and the 2B accessory domain is colored in medium gray. (**D**) AlphaFold prediction of the complex formed between HELB (gray) and trimeric RPA (yellow) predicts interaction of the HSM with the RPA70 N-terminal domain (NTD). The apparent interactions of the RPA core with HELB (left side of D) are low confidence predictions (**Supplementary Figure S2**). (**E**) A crystal structure of a HELB peptide with the RPA70 NTD (PDB:7XV1) shows that E499, D506, and D510 (all magenta) interact with RPA (26).

## MATERIAL AND METHODS

### AlphaFold prediction

The amino acid sequences for HELB (NP_001357214.1), RPA14 (NP_002938.1), RPA32 (NP_002937.1), and RPA70 (NP_002936.1) were supplied to AlphaFold-Multimer, version 2.3.1 (25, 28), one copy of each sequence. The program was setup with default parameters, which generates 5 models with 5 predictions each for a total of 25 multimer structure predictions. The database date cutoff was set to 2023-01-10. The highest-ranking model had a scores of ptm: 0.522, iptm: 0.451, and rank: 0.465. The predicted alignment error generated by AlphaFold for this model was plotted with ChimeraX (28–30), and its PDB coordinate file was evaluated and illustrated with PyMOL (31).

### Plasmids, and oligonucleotides

The pcDNA5/FRT/TO-GFP-HELB plasmid for expression of siRNA resistant GFP-tagged HELB in human cells was a gift from Daniel Durocher (21). The pcDNA5/FRT/TO-SFB-HELB plasmid was created by replacing the GFP sequence with an S-tag, 2X Flag tag, and streptavidin-binding peptide (SFB) (32) by Gibson Assembly. pcDNA5/FRT/TO plasmids encoding D506G HELB and E499A/D506A/D510A (3xA) HELB were created using site-directed mutagenesis. A plasmid expressing the tag only, pcDNA5/FRT/TO-SFB-EV (empty vector), was generated by Gibson Assembly. pSpCas9(BB)-2A-Puro (PX459) was a gift from Feng Zhang (Addgene plasmid # 62988) (33).

Oligonucleotides were ordered from Integrated DNA Technologies; sequences are listed in **Supplementary Table S1**. siGENOME siRNAs were ordered from Horizon Discovery. siGENOME targeting human HELB (D-013541-02: ACAGUCGAACGUUACUUUC) and siGENOME Non-targeting siRNA Pool #1 (D-001206-13) were used.

### Cell culture

HEK293T (ATCC CRL-3216), U2OS (ATCC HTB-96), U2OS-265 (34) and U2OS DR-GFP (ATCC CRL-3455) cells were grown in DMEM (ThermoFisher Scientific 10569044) supplemented with 10% FBS (R&D solutions S11150) at 37°C with 5% CO_2_. U2OS-265 DSB reporter cells were a generous gift from Roger Greenberg (34).

### Generation of HELB knockout cells

CRISPR/Cas9 was used to generate HELB knockout clones in U2OS and HEK293T cell lines as described previously (33). Briefly, two gRNAs targeting exon 1 in all splice variants of HELB were generated using the online tool at www.atum.bio/eCommerce/cas9/input and were synthesized by Integrated DNA Technologies. The sequences are listed in **Supplementary Table S1**. The gRNAs were annealed, phosphorylated by T4 polynucleotide kinase, and ligated into the BsbI digested pSpCas9(BB)-2A-Puro (PX459) plasmid with T7 DNA ligase. Next, exonuclease digestion of residual linear plasmid was conducted by PlasmidSafe Exonuclease. Cells were then transfected with the construct encoding the appropriate gRNA. pSPCas9(HELB gRNA) plasmid (10 μg) was mixed with 50 μl of 1 mg/ml polyethylenimine, PEI (Polysciences, Inc) in 37 °C OptiMEM I by vortexing. Following a 30-minute incubation at room temperature, plasmid mix was added dropwise to cells cultured in antibiotic-free medium DMEM. After 24-hours of transfection, clones were selected for 48 hours with 2 μg/mL of puromycin prior to plating and isolating individual clones. The HELB knockout clones were validated by western blotting (**Supplementary Figure S2**).

### Western blotting

Cells were harvested and resuspended in lysis buffer (40 mM HEPES pH 7.5, 10 mM NaCl, 1% Triton X-100, 1X protease inhibitor cocktail (Sigma P2714), 20 mM β-glycerophosphate, 1 mM Na_2_VO_4,_ and 1 mM dithiothreitol). Proteins were separated by 7.5% SDS-PAGE, and the proteins were transferred to a 0.45 µm nitrocellulose membrane. The membrane was blocked overnight with 5% non-fat milk in 1X TBS with 0.1% Tween-20 (TBS-T). Membranes were incubated with primary antibodies in TBS-T with 1% milk for 1 hour at room temperature. Then membranes were incubated with HRP conjugated secondary antibodies in TBS-T with 1% milk for 1 hour at room temperature. Blots were imaged using Amersham ECL Prime (GE Healthcare) on a ChemiDoc MP imaging system (Bio-Rad). Antibodies used were rabbit anti-HELB (Abcam: ab202141, 1:10,000-1:5,000), rabbit anti-RPA70 (Cell Signaling 2267S, 1:1000), mouse anti-β-actin (Cell signaling: 3700S, 1:2000), mouse anti-FLAG (Sigma F1804, 1:1000), HRP labelled goat anti-mouse IgG (Perkin Elmer: NEF822001EA, 1:10,000), HRP labelled goat anti-rabbit IgG (Perkin Elmer: NEF812001EA, 1:10,000), StarBright Blue 520 goat anti-mouse IgG (Bio-Rad 64456855, 1:2,500), StarBright Blue 700 goat anti-rabbit (Bio-Rad 64484700, 1:2,500).

### Tandem affinity purification mass spectrometry (TAP-MS)

HELB^KO^ HEK293T cells were transfected with plasmids encoding pcDNA5/FRT/TO-SFB-HELB variants or pcDNA5/FRT/TO-SFB-EV (empty vector) using PEI (Polysciences, Inc). After 24 hours, the plates were split, and 48 hours after transfection, the cells were harvested by trypsinization. The cell pellet was lysed in NETN (150 mM NaCl, 0.5 mM EDTA, 20 mM Tris-Cl pH 8,0, 0.5% NP-40 Alternative (Millipore 492016-500)) supplemented with 1 µg/mL aprotinin, and 1 µg/mL pepstatin A with rotation for 1 hour at 4°C. An aliquot of the lysate was removed and saved for normalization. The samples were centrifuged at 9,000x*g* for 20 min at 4°C. The supernatant (soluble fraction) was transferred to a new tube. Streptavidin Sepharose High Performance (Cytiva) was added to the tubes containing the supernatants, and the samples were rotated for 1 hour at 4°C.

While the soluble fraction was rotating, the chromatin pellet was washed with PBS before adding NETN supplemented with 1 µg/mL aprotinin, 1 µg/mL pepstatin A, 10 mM MgCl_2_, and 200 U Turbonuclease from *Serratia marcescens* (Sigma). Samples were rotated for 1 hour at 4°C before centrifuging at 9,000x*g* for 20 minutes at 4°C. The supernatants (chromatin fraction) were transferred to new tubes.

Streptavidin Sepharose High Performance (Cytiva) was added to the tubes containing the supernatants, and the samples were rotated for 1 hour at 4°C. Samples were centrifuged at 211x*g* for 2 minutes before aspirating the supernatant. The pellets were resuspended in NETN and transferred to a clean tube where the pellets were washed three times with NETN. After aspiration of the last wash, SFB-tagged protein complexes were eluted by addition of 2 mg/ml biotin in NETN with rotation for 1 hour at 4°C followed by centrifugation at 211*xg* for 1 minute. The supernatant was transferred to a new tube and the biotin elution was repeated. After the second biotin elution, S protein agarose (Millipore) was added to each sample, and the samples were rotated for 1 hour at 4°C before washing three times with NETN.

To remove detergents and salts for mass spectrometry, the beads were washed in freshly prepared, cold, 100 mM ammonium bicarbonate, transferred to a new tube, and washed a second time. After removing the supernatant, the beads were frozen at −80°C, and submitted to the IDeA National Resource for Quantitative Proteomics for analysis. Experiments were performed in biological quintuplicate.

### Streptavidin pulldown

HELB^KO^ HEK293T cells were transfected with plasmids encoding pcDNA5/FRT/TO-SFB-HELB variants using PEI (Polysciences, Inc). After 24 hours, cells were harvested by trypsinization. The cell pellet was lysed in NETN (150 mM NaCl, 0.5 mM EDTA, 20 mM Tris-Cl pH 8,0, 05% NP-40) supplemented with 1 μg/mL aprotinin, 2 μg/mL pepstatin A, 10 mM MgCl_2_, and 100 U Turbonuclease from *Serratia marcescens* (Sigma). Samples were rotated for 1 hour at 4°C. Samples were centrifuged at 13,000x*g* for 20 minutes at 4°C. The supernatants were transferred to clean tubes, and a sample of the supernatant was saved in a separate tube as the input sample. Streptavidin Sepharose High Performance (Cytiva) was added to the tubes containing the supernatants, and the samples were rotated for 1 hour at 4°C. Samples were centrifuged at 211x*g* for 1 minute before aspirating the supernatant. The pellets were washed three times with NETN with 1 μg/mL aprotinin, 2 μg/mL pepstatin A. After aspiration of the last wash, Laemmli buffer was added to the beads and the reserved input sample. Both were heated at 95°C for 5 min before separation by 7.5% SDS-PAGE and western blotting.

### Immunofluorescence

pcDNA5/FRT/TO-SFB-HELB (WT, D506G, and 3xA) plasmids (1 µg) were transfected into 1.0 × 10^5^ cells using FuGENE HD (Promega) according to the manufacturer’s protocol followed by incubation at 37°C with 5% CO_2_ 24 hours. After incubation, cells were treated 10 µM camptothecin (CPT) for 1 hour at 37°C with 5% CO_2_. Post treatment, cells were washed with 1X PBS before pre-extraction with 0.5% Triton X-100 for 5 minutes on ice. Cells were washed again with 1X PBS and fixed with 2% paraformaldehyde for 15 minutes at room temperature, washed again with 1X PBS and permeabilized with a solution consisting of 20 mM HEPES, 50 mM NaCl, 3 mM MgCl_2_, 300 mM sucrose, 0.5% Triton X-100 in Milli-Q water for 10 minutes on ice before blocking overnight with 5% BSA in 1x PBS. Cells were incubated with rabbit anti-FLAG antibody (Cell Signaling: 2368S, 1:1000) for 1 hour at room temperature followed by goat anti-rabbit IgG Alexa Fluor 647 (Invitrogen: A21244, 1:2500) for 1 hour at room temperature. After washing, the cells were mounted with 2.5 mg/ml DABCO (1,4-diazobicyclo[2.2.2]octane) and 0.1 μg/ml DAPI in 25% PBS, 75% glycerol. Cells were imaged on an Olympus FV1000 confocal microscope. The intensity of HELB in the nucleus of each cell was quantified using CellProfiler (35). Experiments were performed in triplicate.

### FokI nuclease double-strand DNA break co-localization assay

U2OS-265 cells expressing a DSB reporter were a gift from Roger Greenberg (34). Cells were seeded in a 6-well dish before transfecting with plasmids encoding GFP-HELB (WT, D506G, or 3xA). The next day, cells were moved to glass coverslips. After 24 hours, cells were treated with 1 μM 4-hydroxytamoxifen (4-OHT) (Sigma) and 1 μM Shield1 (Clonetech Labs) for 4 hours. Cells were fixed using 3% paraformaldehyde, permeabilized, blocked for 1 hour with 5% BSA, and stained with rabbit anti-mCherry (Cell Signaling 43590S, 1:200) followed by goat anti-rabbit IgG Alexa Fluor 594 (Invitrogen: A11037, 1:500) and Hoechst 33342 (Invitrogen H3570, 1:10,000) before mounting with anti-fade solution (0.02% p-phenylenediamine [Sigma, P6001] in 90% glycerol in PBS). Cells were imaged on a Nikon Eclipse Ti2 confocal microscope. Experiments were performed in triplicate, and localization of GFP-HELB relative to mCherry was scored blind.

### DR-GFP HR reporter assay

DR-GFP U2OS cells were transfected with 89.6 pmol of siGENOME non-targeting pool and siGENOME human HELB (Horizon Discovery) using DharmaFECT 1 (Horizon Discovery). DharmaFECT 1 and siRNA were separately diluted in OptiMEM before combining and incubating 20 minutes at room temperature. After addition of the transfection agent to the cells, they were incubated at 37°C with 5% CO_2_. 24 hours after siRNA transfection, the media with transfection reagent was replaced with a fresh media. Cells were transfected again with 1.8 µg of plasmid either encoding pcDNA5/FRT/TO-SFB-HELB variants or pcDNA5/FRT/TO-SFB-EV (empty vector) with or without pCBA-iSceI using PEI. Plasmids were mixed with OptiMEM before addition of PEI. After incubation at room temperature for 20-30 minutes, the transfection reaction was added to the cells gently and incubated at 37 °C with 5% CO_2_. 24 hours after the second transfection, the media with transfection was replaced with fresh media and incubated again.

72 hours after siRNA transfection, cells were harvested by trypsinization followed by mixing 2:1 with 10 % formaldehyde and vortexing immediately for 2-3 seconds on medium speed. This was followed by incubation in fixative for 30 minutes. Cells were centrifuged at 150×*g* for 5 min to pellet the cells, fixative was removed, and cells were resuspended in 500 μL PBS. The resuspended cells were transferred into round-bottom tubes with strainer snap caps and centrifuged again at 150x*g* for 5 minutes to further separate cells. Cells were analyzed for green fluorescence on a BD LSRFortessa. Cells were gated by forward scatter versus side scatter. Experiments were performed in biological triplicate. **Supplementary Figure S3C** shows the knockdown efficiency.

## RESULTS

### D506G substitution in HELB impairs interaction with RPA

Because the D506 is one of three residues of HELB reported to interact with RPA (magenta in **Figure 1** and **Supplementary Figure S1**) (19), we investigated whether a D506G substitution affected HELB interactions with other proteins. The protein interactomes of WT, D506G, and 3xA SFB-HELB in the chromatin and soluble fractions were identified using TAP-MS and compared to that of SFB-EV (**Figure 2**, **Supplementary Figure S4**, and **Supplementary Figure S5**). In the WT HELB chromatin fraction, all three subunits of RPA are highly enriched (**Figure 2A**). The RPA70 and RPA14 subunits are also enriched in the D506G HELB chromatin samples (**Supplementary Figure S4A**). However, the degree of enrichment is reduced in the D506G HELB samples relative to WT HELB samples, and all three subunits of RPA are depleted in the D506G HELB relative to WT HELB (**Figure 2B**). This suggests that a D5506G substitution in HELB interferes with interaction with RPA.

**Figure 2.**
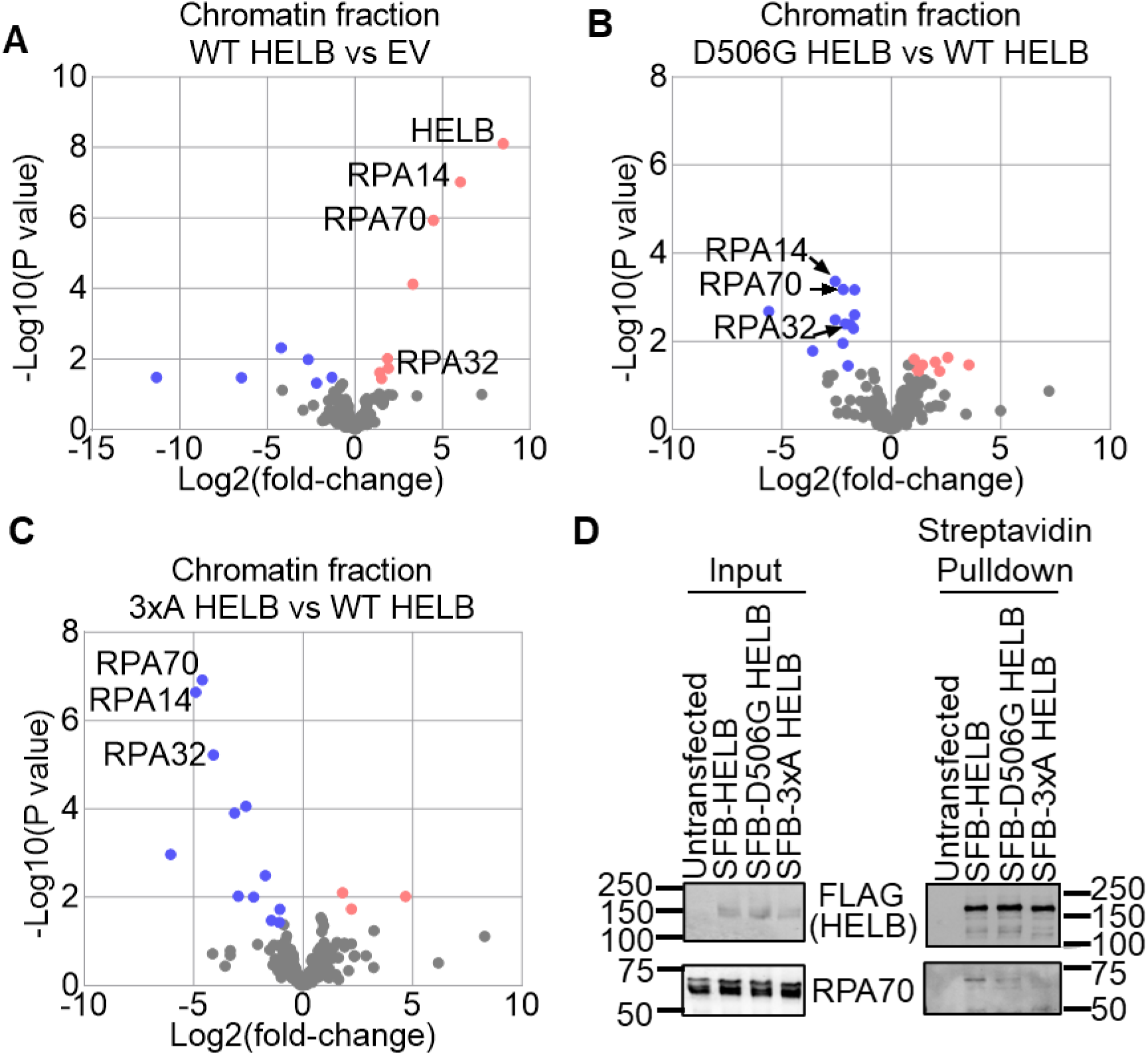
D506G substitution in HELB reduces interaction with RPA. Proteins were identified by TAP-MS from the chromatin fraction isolated from HELB^KO^ 293T cells expressing SFB-EV or SFB-HELB (WT, D506G, or 3xA) in biological quintuplicate. Significantly enriched (red) and depleted (blue) proteins are plotted for WT HELB relative to EV (**A**), D506G HELB relative to WT HELB (**B**), and 3xA HELB relative to WT HELB (**C**). (**D**) Input samples and samples after pulldown of SFB-HELB on streptavidin beads were probed for FLAG and RPA70 by western blot.

None of the subunits of RPA are significantly enriched in the 3xA HELB chromatin samples (**Supplementary Figure S4B**), and they are significantly depleted in the 3xA HELB chromatin samples relative to the WT HELB chromatin samples (**Figure 2C**), confirming the results from the Fanning lab that (19) that E499, E506, and D510 are critical residues for interaction with RPA. Interestingly, very few proteins are significantly enriched in the 3xA HELB samples (**Supplementary Figure S4B**), suggesting that many of the HELB interacting proteins in the chromatin fraction may interact indirectly through RPA.

HELB binds to chromatin with DNA damage and replication stress (19, 21, 23) but is also present in the nucleoplasm and cytoplasm (19). Thus, we also identified HELB interacting proteins in the soluble fraction that contains both the nucleoplasm and cytoplasm (**Supplementary Figure S5**). Besides HELB itself, the three subunits of RPA are the most significant highly enriched proteins in the WT HELB soluble samples (**Supplementary Figure S5A**), suggesting that HELB may interact with RPA in a DNA independent manner. This is consistent with structural studies of a peptide containing a portion of the HSM including E499, E506, and D510 interacting with the RPA70 NTD in the absence of DNA (19, 26).

In the D506G HELB soluble samples, the three subunits of RPA are also enriched, although to a lesser degree than in the WT HELB soluble samples (**Supplementary Figure S5B**), and all three subunits are significantly depleted in the D506G HELB soluble samples relative to the WT HELB soluble samples (**Supplementary Figure S5C**). Similar to in the chromatin fractions, none of the RPA subunits are significantly enriched in the 3xA HELB soluble samples (**Supplementary Figure S5D**) and they are significantly depleted in the 3xA HELB soluble samples relative to the WT HELB soluble samples (**Supplementary Figure S5E**).

In WT HELB, D506G HELB, and 3xA HELB soluble samples, CDK2 was significantly enriched (**Supplementary Figure S5A, B, & D**). CDK2 phosphorylates HELB at the G1 to S transition (23), and this interaction with WT HELB was previously observed in cells treated with neocarzinostatin, a radiomimetic drug (21). This interaction was only observed in the soluble fraction, suggesting that CDK2 phosphorylation occurs when HELB is not bound to chromatin. In addition, because CDK2 was similarly enriched in the WT HELB, D506G HELB, and 3xA HELB soluble samples, this indicates that HELB interaction with CDK2 is independent of interaction with RPA.

Because D506G substitution interfered with HELB interaction with RPA we used affinity purification to confirm the reduction in this important interaction. RPA70 co-purifies with SFB-tagged wildtype HELB (**Figure 2D**), and co-purification of RPA70 with the RPA-interaction deficient SFB-3xA HELB is eliminated (**Figure 2D**). Co-purification of RPA70 with SFB-D506G HELB is greatly reduced (**Figure 2D**), indicating that substitution of D506 alone is sufficient to interfere with HELB interaction with RPA and the HELB encoded by the rs75770066 missense allele has impaired interaction with RPA.

### D506G substitution in HELB interferes with HELB localization to chromatin

HELB localizes to foci on chromatin during DNA replication stress and DNA damage (19, 23, 36). Localization of HELB to sites of laser induced microirradiation is dependent on its interaction with RPA as 3xA HELB is severely impaired in recruitment to sites of microirradiation (21). The necessity of interaction with RPA for recruitment of HELB to other types of DNA damage is unknown. HELB localization to chromatin was measured in HELB^KO^ U2OS cells expressing SFB-tagged HELB variants treated with camptothecin (CPT), a topoisomerase 1 inhibitor that induces DNA replication stress (**Figure 3A-B**). Wildtype SFB-HELB bound to chromatin increases with CPT treatment while SFB-D506G HELB bound to chromatin does not increase. A very small increase in SFB-3xA HELB is observed with CPT treatment, but the level of chromatin-bound HELB is still below that of untreated cells expressing wildtype HELB. This indicates that interaction with RPA is required for localization of HELB to DNA with replication stress and that the missense variant encoded by rs75770066 does not localize to DNA with DNA replication stress.

**Figure 3.**
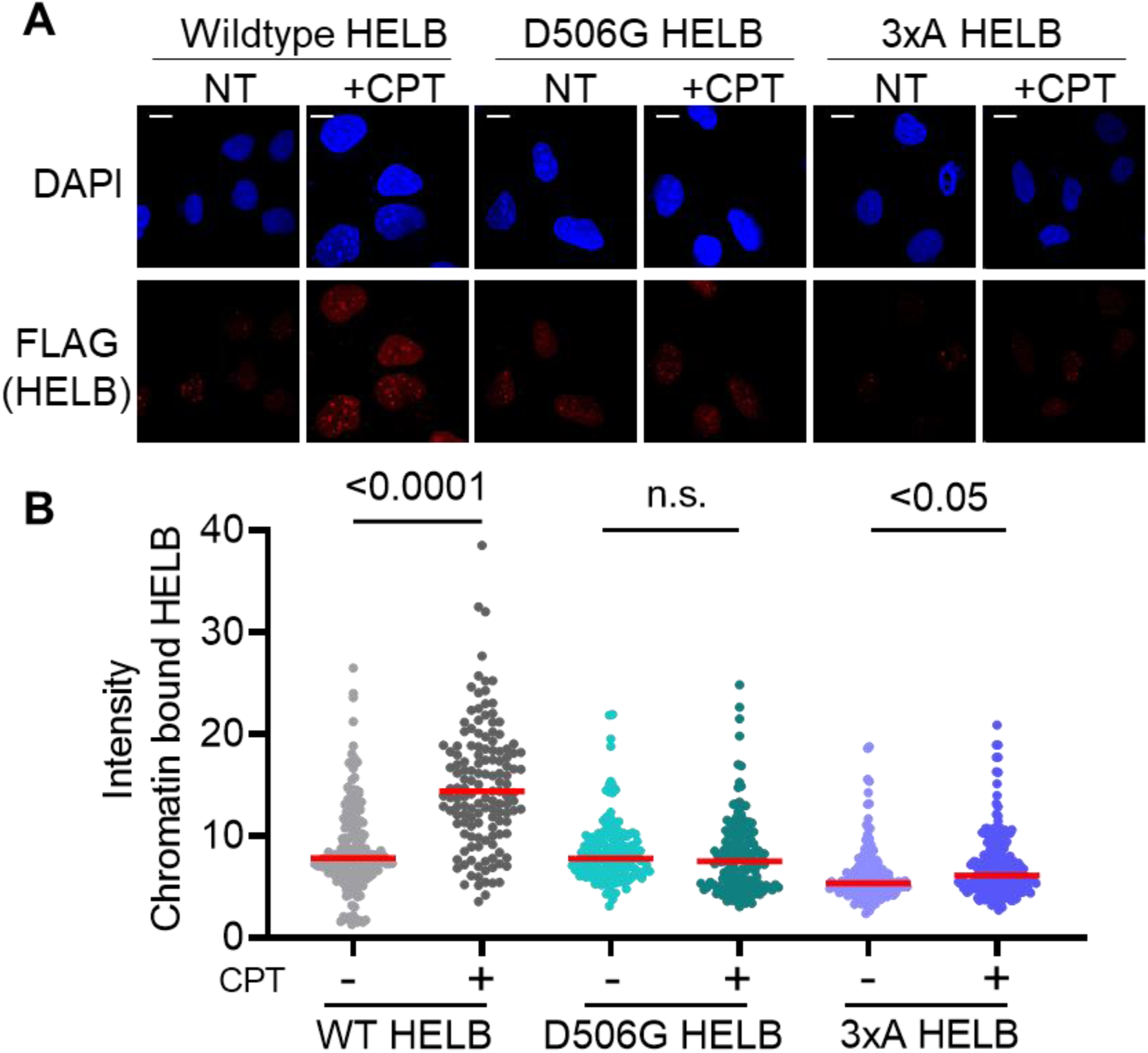
D506G substitution in HELB interferes with localization of HELB to chromatin in response to replication stress. (**A**) HELB^KO^ U2OS cells expressing SFB-HELB variants (WT, D506G, and 3xA) were treated with 10 μM camptothecin (CPT) for 1 hour followed by pre-extraction to remove non-chromatin-bound proteins before fixing and staining with an anti-FLAG antibody to detect SFB-HELB. (**B**) The intensity of HELB in the nuclei of >130 cells for each condition was quantified using CellProfiler (35) from biological triplicate experiments. Scale bar indicates 1 μm.

We also measured the ability of HELB variants to localize to double stranded DNA breaks (DSBs) using a DSB reporter (34). The DSB reporter cells stably express an mCherry-LacI-FokI fusion protein that creates DSBs at a single locus on chromosome 1 containing 256 copies of the lac operator (37). The fusion protein also expresses a modified estradiol receptor for nuclear targeting and a destabilization domain to prevent unwanted DSBs (34). Following treatment of DSB reporter cells with 4-hydroxytamoxifen (4-OHT) and Shield1, the fusion protein is stabilized and DSBs are induced (**Figure 4A**). GFP-tagged HELB variants were expressed in the DSB reporter cells and co-localization of mCherry-FokI and GFB-HELB was measured. Approximately 60% of the mCherry-FokI foci co-localize with wildtype GFP-HELB (**Figure 4B-C**). The lack of co-localization of GFP-HELB with mCherry-FokI in all cells could be due to the cell cycle dependent localization of HELB (21, 23). Co-localization of both GFP-D506G HELB and GFP-3xA HELB is reduced to approximately 30% (**Figure 4B-C**). A corresponding increase in the fraction of mCherry-FokI foci that do not co-localize with HELB is observed in cells expressing GFP-D506G HELB and GFP-3xA HELB (**Figure 4B-C**). This indicates that interaction with RPA promotes HELB localization to DSBs. In addition, recruitment of the D506G variant is reduced relative to wildtype HELB.

**Figure 4.**
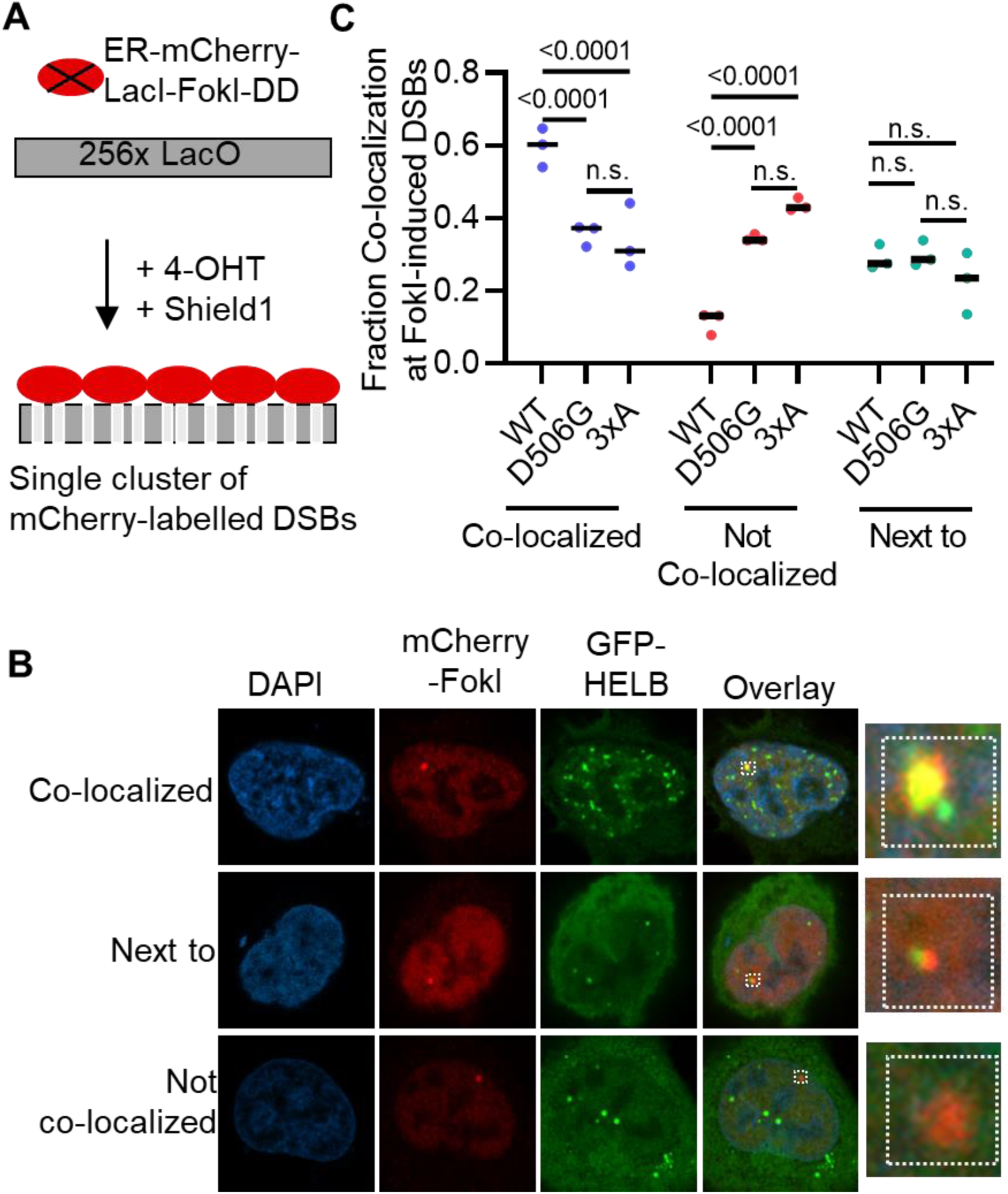
D506G HELB has reduced co-localization with DSBs relative to WT HELB. (**A**) A single array of mCherry labeled DSBs is produced upon treatment of DSB reporter cells with 4-hydroxytamixofen (4-OHT) and Shield1. (**B**) DSB reporter cells expressing GFP-HELB variants (WT, D506G, and 3xA) were treated with 1 μM 4-OHT and 1 μM Shield1 for 4 hours. Cells were fixed and stained with an antibody to mCherry before imaging. (**C**) Localization of GFP-HELB relative to Cherry-FokI foci was quantified blind for >150 cells per condition from biological triplicate experiments.

Interestingly, in each of the conditions, ∼25% of the mCherry-FokI foci have GFP-HELB localized immediately next to but not overlapping with the mCherry-Fok1 (**Figure 4B-C**). DSBs induce formation of a new chromatin compartment (38), and at least three separate subcompartments have been observed at DSBs, one containing ssDNA and end resection factors, another containing HR proteins, and a third containing proteins involved in non-homologous end joining (39). It is possible that HELB is present in two different DNA repair compartments, one co-localized with the DSB and one localized immediately adjacent to the DSB. Because similar quantities of GFP-HELB foci next to mCherry-FokI foci are observed with WT, D506G, and 3xA HELB, this suggests that localization of HELB adjacent to DSBs is not dependent on interaction with RPA. It is possible that some of the cells where GFP-HELB appears to be co-localized with mCherry-FokI in **Figure 4** may actually be foci where GFP-HELB is located next to the mCherry-FokI focus but appear to be co-localized due to the angle of the image. Because the fraction of cells with GFP-HELB localized next to the mCherry-FokI focus is the same for WT HELB, D506G HELB, and 3xA HELB, this is unlikely to affect the relative amount of GFP-HELB co-localized with mCherry-FokI in the different samples but could inflate the absolute levels of GFP-HELB co-localized with mCherry-FokI in each of the samples.

### D506G substitution in HELB increases repair of DSBs by HR

The impaired localization of D506G HELB to DSBs suggests regulation of HR by HELB (21, 36) may also be impaired. We measured recombination in U2OS cells containing a DR-GFP reporter (40). This reporter contains a direct repeat of two *GFP* genes: *I-SceI GFP* contains the recognition site for I-SceI endonuclease and two stop codons and *iGFP* is an internal fragment of the *GFP* gene that can be used as a repair template during HR, restoring the *GFP* gene (**Figure 5A**). GFP positive cells were absent in cells not transfected with a plasmid encoding I-SceI (**Figure 5B**). Transfection with plasmids encoding I-SceI and SFB-empty vector results in ∼1.2% GFP positive cells. Consistent with previous results (21), we observed a 50% reduction in GFP positive cells when transfecting with plasmids encoding I-SceI and wildtype HELB (**Figure 5B**). Similarly, we saw an increase in GFP positive cells when we treated with siRNA targeting HELB and transfected with plasmids encoding I-SceI and empty vector. No increase in GFP positive cells was observed when cells were transfected with a plasmid encoding wildtype HELB that rescued knockdown of HELB expression. However, neither expression of D506G HELB or 3xA HELB could rescue knockdown of HELB expression; in both cases, recombination rates were increased similarly to those of cells with knockdown of HELB expression and expression of an empty vector. Therefore, interaction with RPA is required for HELB to limit HR, and the D506G missense variant caused by rs75770066 interferes with HELB’s ability to inhibit HR.

**Figure 5.**
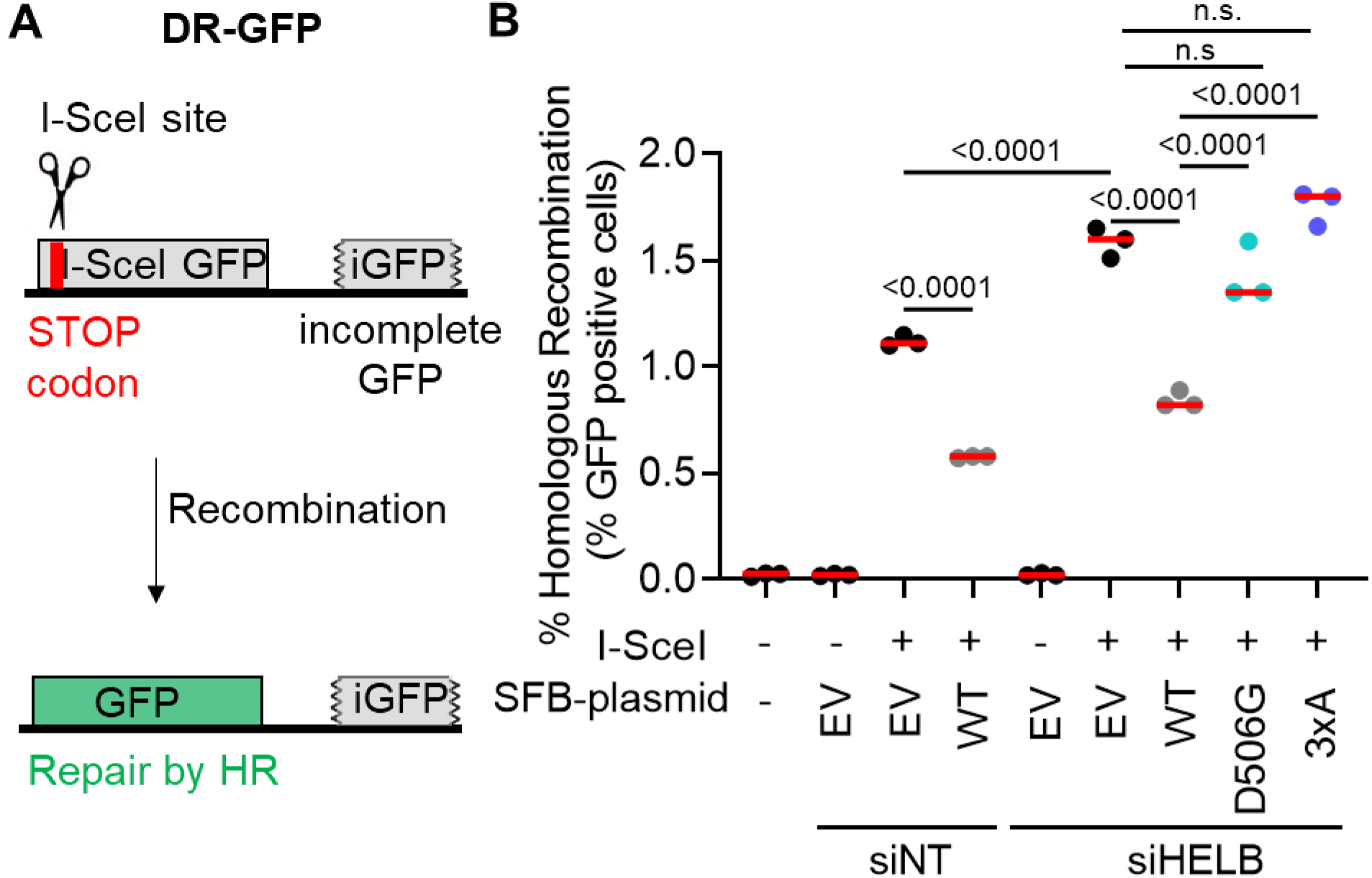
Expression of D506G HELB increases HR levels. (**A**) Expression of I-SceI in DR-GFP-U2OS cells produces a DSB in the I-SceI GFP gene. Cells that repair the DSB by HR are identified as GFP-positive cells by flow cytometry. (**B**) The percentage of GFP-positive cells in each sample are plotted.

## DISCUSSION

There are multiple SNPs in *HELB* that are associated with altered ANM in women. In addition, multiple SNPs in *HELB* are associated with reproductive traits in male and female cattle (41, 42), suggesting HELB plays a critical role in reproductive fitness. In humans, rs75770066 is a low frequency variant resulting in a D506G substitution in the protein that alters ANM (6, 12–14). We compared the interaction partners and cellular activity of D506G HELB to WT HELB to gain insight into how rs75770066 may alter ANM. D506 is located in the HSM insertion within the helicase domain of HELB (**Figure 1A-C**). The HSM partially overlaps with an acidic motif previously described in HELB (19) and contains three negatively charged amino acids (E499, D506, and D510) previously shown to interact with RPA (19, 21, 26). Consistent with this prior data, AlphaFold predicts that the HSM interacts with the basic cleft on the RPA70 NTD (**Figure 1D**). The HSM is an insertion within the 1A domain of HELB. In SF1B helicases like HELB that unwind DNA with a 5′-to-3′ polarity, the 1A domain is the leading edge of the enzyme (43–48). This positions the RPA interacting region in front of HELB, which is consistent with the ability of HELB to clear RPA from ssDNA (22). We found that WT HELB interacts with RPA in both the chromatin and soluble fractions (**Figure 2**, **Supplementary Figure S4**, **Supplementary Figure S5**), suggesting that HELB may not require DNA to interact with RPA. This possibility is consistent with structural data indicating interaction of a peptide of HELB including E499, E506, and D510 with the RPA70 NTD in the absence of DNA (19, 26). D506G substitution reduces interaction of HELB with several proteins including all three subunits of RPA (**Figure 2**), but does not affect interaction with CDK2 (**Supplementary Figure S5**).

In cells, HELB performs several important functions that may affect cell growth and viability. HELB promotes timely entry into S phase (18, 23), enhances recovery from DNA replication stress (19), and regulates HR (21, 36). Recruitment of HELB to sites of laser induced microirradiation has previously been shown to be dependent on interaction with RPA (21). In cells expressing RPA interaction deficient 3xA HELB, we found that recruitment of HELB to chromatin in response to replication stress (**Figure 3**) and recruitment to DSBs (**Figure 4**) are also dependent on interaction with RPA. In addition, we found that interaction with RPA is required for HELB to inhibit HR (**Figure 5**). In their previous report that HELB inhibits HR, the Durocher group showed that HELB reduces long range end-resection by EXO1 and BLM-DNA2 (21). They found that inhibition of end-resection required interaction with RPA as cells expressing the 3xA HELB variant had similar levels of ssDNA produced by resection as cells lacking HELB (21), suggesting that interaction with RPA is required for HELB to inhibit HR. The results in **Figure 5** confirm that this is indeed the case.

D506G substitution also interferes with the recruitment of HELB to chromatin with DNA replication stress (**Figure 3**) and to sites of DSBs (**Figure 4**) likely because D506G substitution reduces HELB interaction with RPA. Because D506G interferes with HELB recruitment to DSBs, it is not surprising that D506G HELB is unable to suppress HR at DSBs. HR levels in the cells expressing D506G HELB are the same as in cells with expression of HELB knocked down with siRNA or in cells expressing 3xA HELB (**Figure 5**). Thus, D506G substitution in HELB results in impaired cellular function.

Interestingly, HELB has been reported to both stimulate (36) and inhibit (21) HR at DSBs. One group (36) measured HR in SW480/SN.3 colon carcinoma cells containing a SCneo recombination reporter (49). The other group (21) measured HR in HeLa cervical carcinoma cells containing a DR-GFP recombination reporter (40). Although the two groups used different reporter assays, both are based on repair of an I-SceI induced DSB by HR, suggesting the different outcomes are not likely due to the different reporter assays. The group that observed inhibition of HR by HELB (21) transfected with siRNA targeting HELB 24 hours prior to transfection with a plasmid encoding I-SceI while the group that observed HELB enhanced HR (36) transfected with shRNA targeting HELB and a plasmid encoding I-SceI simultaneously. Although both groups observed knockdown of HELB expression 48 hours after transfection with plasmid, it is unlikely that knockdown of HELB expression occurred on the same timescale as I-SceI production in the experiments that observed HELB enhanced HR (36). However, this is also not the likely cause of the contrasting results because insufficient knockdown of HELB expression would likely have no effect on recombination levels instead of producing the opposite effect. It is possible that the different cell lines are the cause of the disparate results. We measured HR levels in a third human cell line, U2OS osteosarcoma cells containing the DR-GFP reporter and observed inhibition of HR by HELB and a corresponding increase in HR in cells depleted of HELB, as was observed by (21). Thus, in at least two human cell lines. HELB is a negative regulator of HR.

Because natural menopause occurs when the oocyte pool is depleted below the threshold necessary for proper ovarian function, changes in the size of the oocyte pool due to decreased survival with replication stress or altered recombination levels may lead to changes in ANM due to impaired establishment of the ovarian reserve. The size of the ovarian reserve is fixed soon after birth by clearing of defective oocytes during the perinatal period (50, 51) (**Figure 6A**). Among the reported cellular functions of HELB, only an increase in recombination levels caused by D506G HELB has the potential to affect oocyte viability negatively or positively (**Figure 6B**). Repair of programed meiotic DSBs by recombination is essential for chromosome pairing via crossover formation and proper segregation in meiosis I. Because D506G HELB is unable to inhibit HR, one allele encoding D506G HELB may enhance recombination and increase oocyte viability. However, a second D506G allele may increase recombination to the point that genomic rearrangements occur, leading to diminished ovarian reserves and early ANM. Interestingly, both insufficient crossover formation and excess crossover formation lead to impaired chromosome segregation, which results in aneuploidy (52, 53). In addition, excessive recombination causes non-allelic homologous recombination, resulting in genomic rearrangements (54). Because both aneuploidy and non-allelic homologous recombination reduce the viability of the oocyte, either too much recombination or too little recombination will reduce the size of the oocyte pool. Failure to correctly repair meiotic DSBs by recombination activates the checkpoint kinases, CHK1 and CHK2, leading to apoptosis (50, 55). In humans, less than 10% of oocytes that begin meiosis survive to puberty (56). This suggests that failure to correctly repair meiotic DSBs by recombination would reduce the size of the ovarian reserve, leading to premature menopause. Thus, the inability of D506G HELB to regulate HR may result in altered ANM caused by rs75770066.

**Figure 6.**
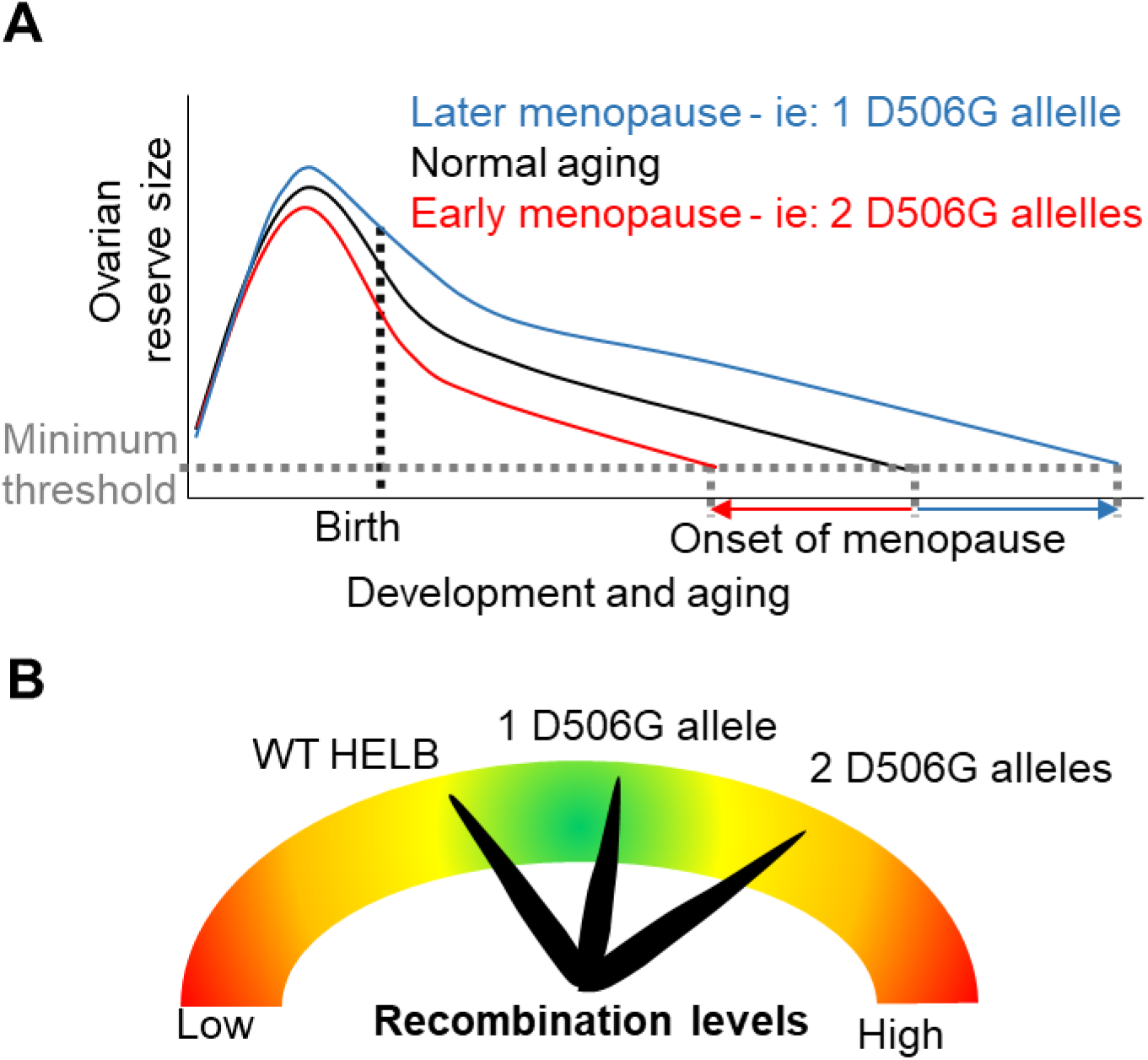
Model for HELB effect on age at menopause. (A). Menopause occurs when oocyte levels drop below a minimum threshold. A single D506G allele may increase the size of the ovarian reserve resulting in delayed ANM while two D506G alleles may decrease the size of the ovarian reserve resulting in early ANM. (B) Repair of DSBs by recombination is essential for pairing and segregation of homologous chromosomes during meiosis. One allele encoding D506G HELB may enhance recombination and increase oocyte viability, but a second D506G allele may increase recombination enough that genomic rearrangements occur, leading to diminished ovarian reserves and early ANM.

We found that D506G substitution in HELB significantly impairs HELB interaction with RPA which interferes with HELB recruitment to sites of DNA damage, including DSBs. D506G HELB is unable to regulate repair of DSBs by HR which may affect oocyte viability, thereby altering ANM.

## SUPPLEMENTARY DATA

In accompanying document

## AUTHOR CONTRIBUTIONS

AKB conceived the study; BO, BHM, CMS, KLW, MKZ, MDT, EB, EJE, and AKB performed the experiments; BO, BHM, CMS, KLW, JLW, EJE, and AKB contributed to data analysis and interpretation; AKB wrote the manuscript with input from all of the authors.

## ACKNOWLEDGEMENTS

We thank Daniel Durocher and Feng Zhang for the kind gifts of plasmids and Roger Greenberg for the kind gift of cells. We thank Dr. Mari Davidson for careful reading of the manuscript and helpful suggestion.

