## Supplementary for "Rare SNP in the *HELB* gene interferes with RPA interaction and cellular function of HELB"

**Supplementary Table S1. gRNA sequences**

| Name | Sequence* |
|---|---|
| HELB gRNA 1 | GGATTTGCTGCTGCACCTCCTCC |
| HELB gRNA 2 | GGAGCCGCAGCTACTTGTGCCTG |

*PAM sequences are underlined.

**Supplementary Figure S1**

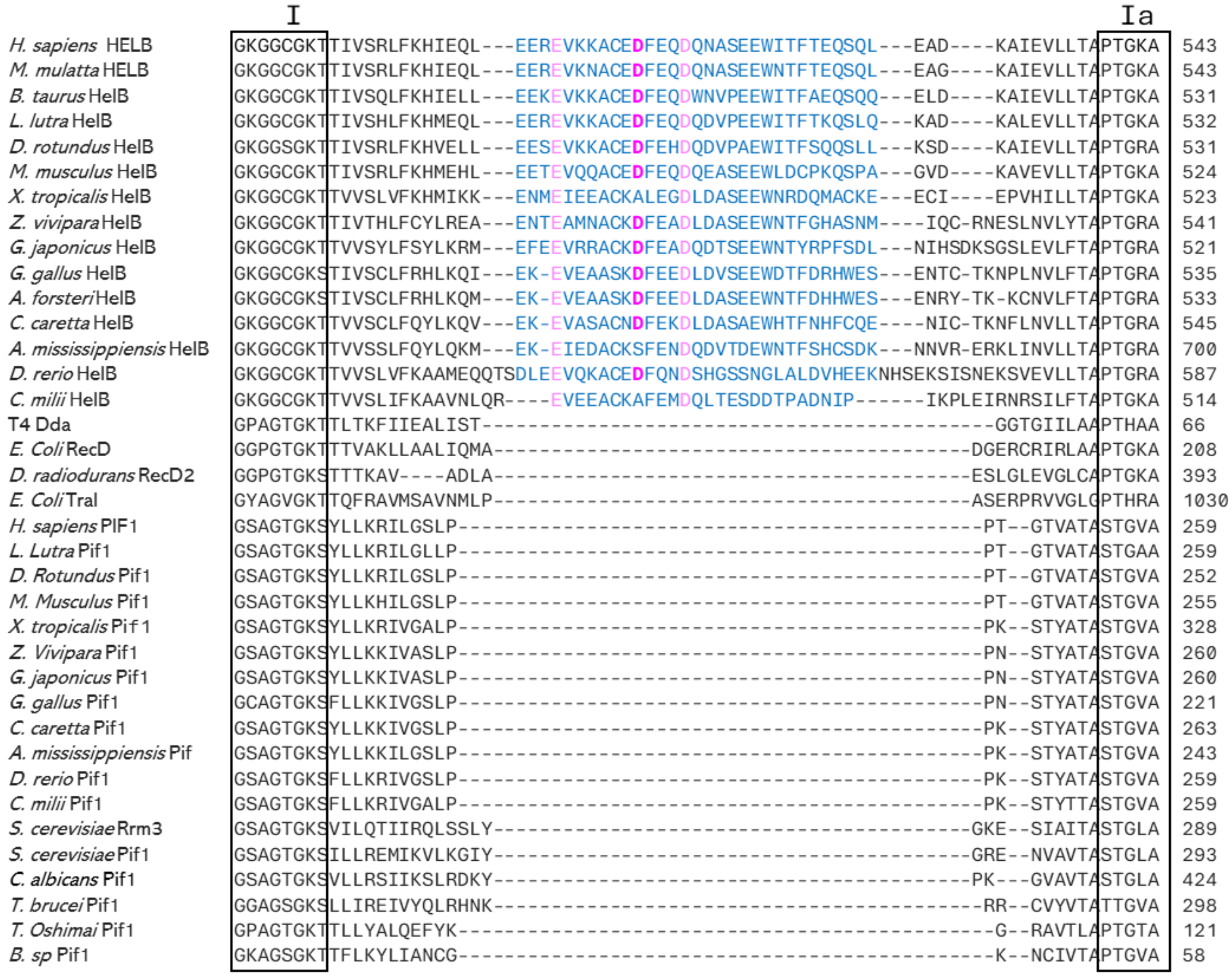


**Supplemental Figure S1**. HELB family proteins contain a HELB specific motif (HSM). Sequence alignments of *Homo sapiens* HELB, *Macaca mulatta* HELB, *Bos taurus* HelB, *Lutra lutra* HelB, *Desmondus rotundus* HelB, *Mus musculus* HelB, *Xenopus tropicalis* HelB, *Zootoca vivipara* HelB, *Gekko japonicus* HelB, *Gallus gallus* HelB, *Aptenodytes forsteri* HelB, *Caretta caretta* HelB, *Alligator mississippiensis* HelB, *Danio rerio* HelB, *Callorhinchus milii* HelB, Bacteriophage T4 Dda, *E. coli* RecD, *Deinococcus radiodurans* RecD2, *E. coli* TraI, *Homo sapiens* PIF1, *Lutra lutra* Pif1, *Desmondus rotundus* Pif1, *Mus musculus* Pif1, *Xenopus tropicalis* Pif1, *Zootoca vivipara* Pif1, *Gekko japonicus* Pif1, *Gallus gallus* Pif1, *Caretta caretta* Pif1, *Alligator mississippiensis* Pif1, *Danio rerio* Pif1, *Callorhinchus milii* Pif1, *Saccharomyces cerevisiae* Rrm3, *Saccharomyces cerevisiae* Pif1, *Candida albicans* Pif1, *Trypanosoma brucei* Pif1, *Thermus oshimai* Pif1, and *Bacteroides sp* Pif1 helicases by Clustal Omega shows a HELB specific motif (HSM) (blue) exists between helicase motifs I and Ia. E499, D506, and D510 (magenta) are located within this HSM.

**Supplementary Figure S2**

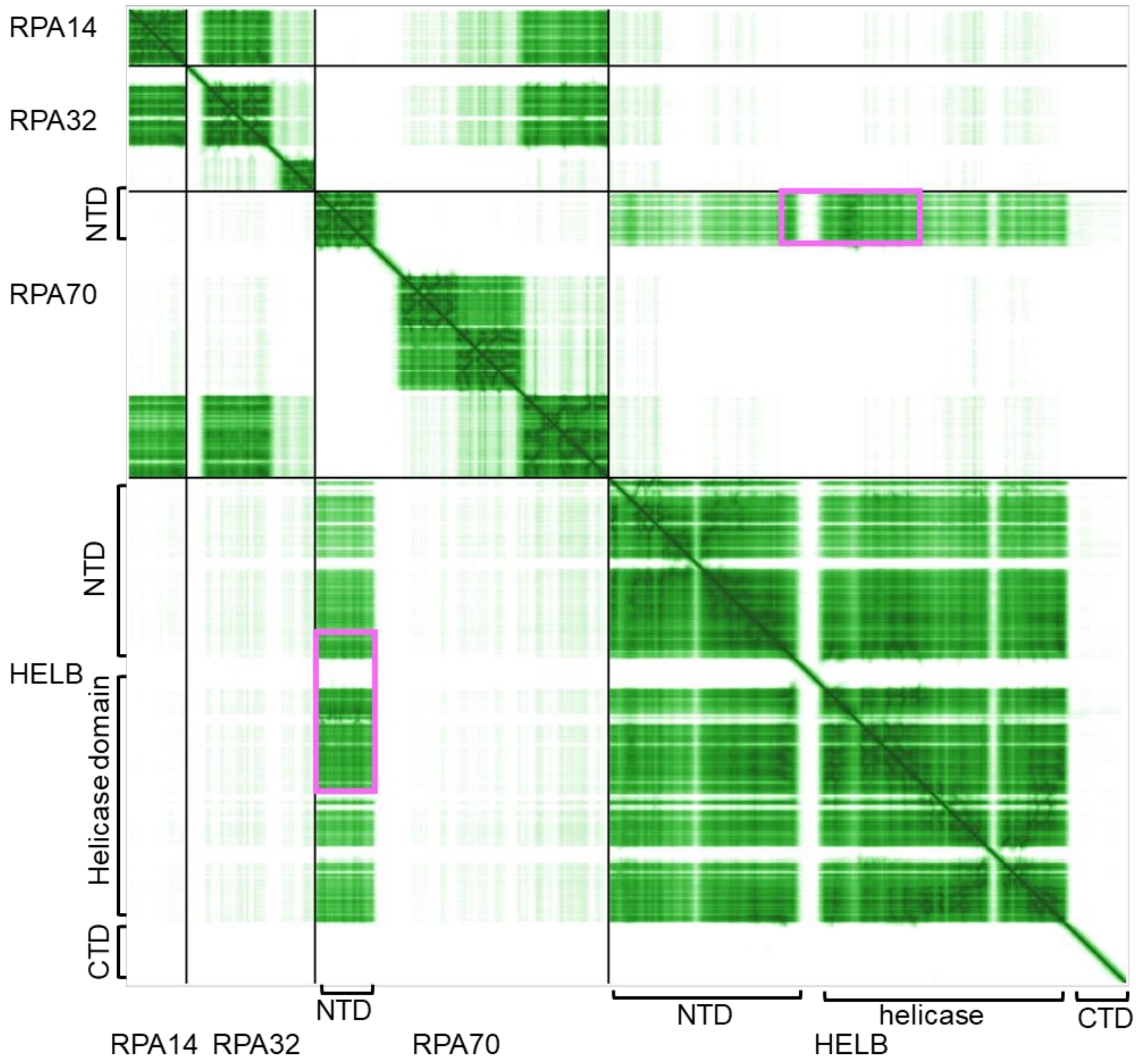


**Supplemental Figure S2**. Only the interaction between the RPA70 NTD and HELB is predicted with confidence by AlphaFold (1, 2). The predicted aligned error for alignment of RPA14, RPA32, RPA70, and HELB is shown with the darker green indicating lower predicted error. Interactions between the RPA70 NTD and the HELB NTD and helicase domains are predicted (pink).

**Supplementary Figure S3**

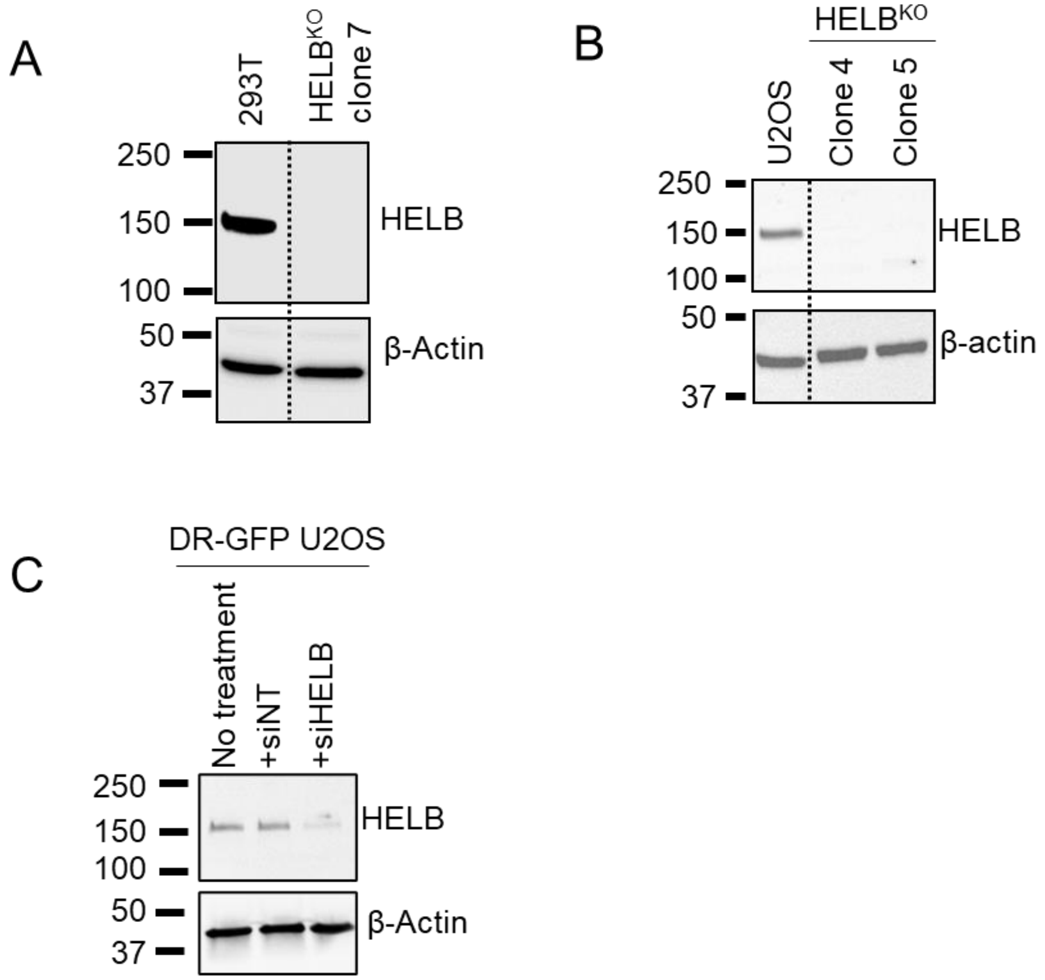


**Supplementary Figure S3**. Western blots illustrating knockout of HELB and knockdown of HELB expression (**A**) Western blot of 293T cells and HELB$^{KO}$ 293T cells. (**B**) Western blot of U2OS cells and HELB$^{KO}$ U2OS cells. (**C**) Western blot of DR-GFP-U2OS cells treated with non-targeting siRNA (siNT) or siRNA against HELB (siHELB).

**Supplementary Figure S4**

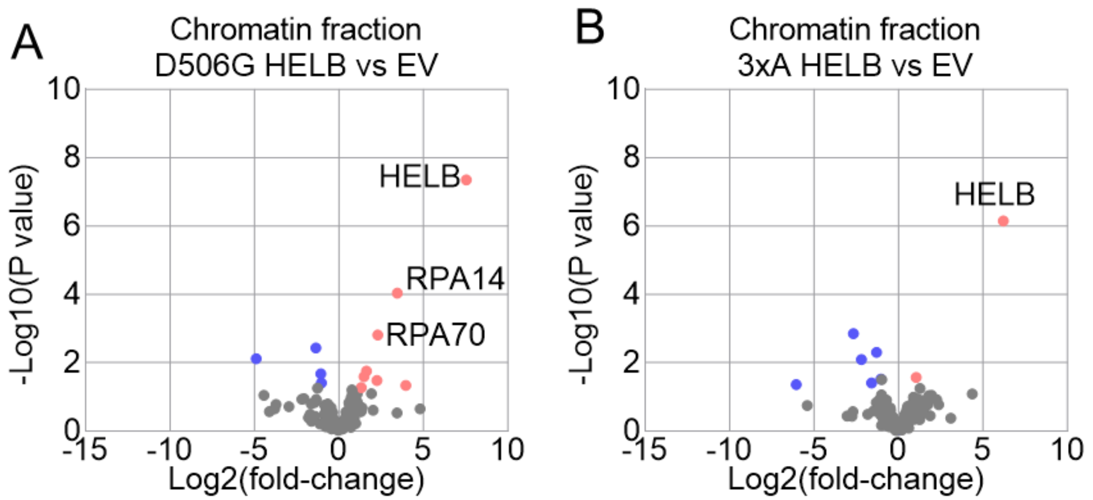


**Supplementary Figure S4**. D506G HELB and 3xA HELB interacting proteins in the chromatin fraction. Proteins were identified by TAP-MS from the chromatin fraction isolated from $HELB^{KO}$ 293T cells expressing SFB-EV or SFB-HELB (D506G, or 3xA). Significantly enriched (red) and depleted (blue) proteins are plotted for D506G HELB relative to EV (**A**) and 3xA HELB relative to EV (**B**).

**Supplementary Figure S5**

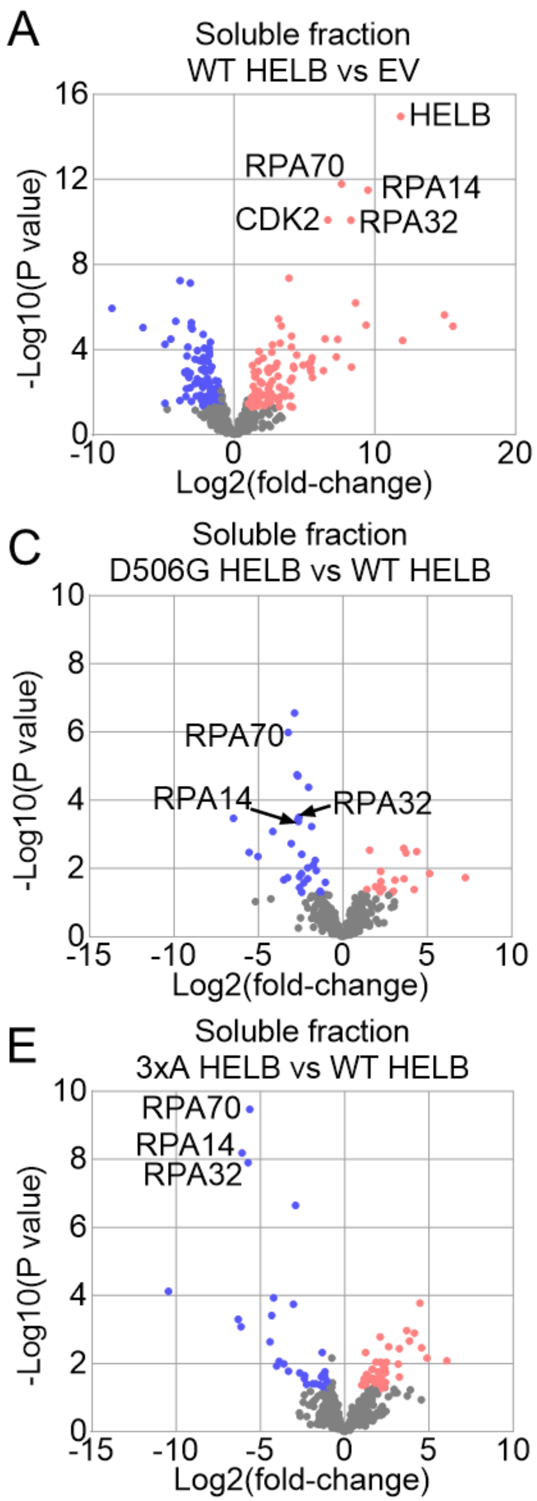


**Supplementary Figure S5**. D506G HELB and 3xA HELB have reduced interactions with RPA in the soluble fraction relative to WT HELB. Proteins were identified by TAP-MS from soluble fractions isolated from HELB$^{KO}$ 293T cells expressing SFB-EV or SFB-HELB (WT, D506G, or 3xA). Significantly enriched (red) and depleted (blue) proteins are plotted for WT HELB relative to EV (**A**), D506G HELB relative to EV (**B**), D506G HELB relative to WT HELB (**C**), 3xA HELB relative to EV (**D**), and 3xA HELB relative to WT HELB (**E**).
